# Strong leaders promote cooperation in heterogeneous populations

**DOI:** 10.64898/2026.07.09.737424

**Authors:** Carlo Longhi, Luis A. Martinez-Vaquero, Fernando Wario Vázquez, Vito Trianni

## Abstract

Many proposed mechanisms for the evolution of cooperation among unrelated individuals rely on relatively demanding cognitive abilities that are not widespread across taxa. In contrast, individual heterogeneity is a pervasive feature of animal groups, encompassing differences in personality as well as physical and cognitive traits. Such heterogeneity can promote the evolution of cooperation, yet its role has received comparatively little attention, particularly as a source of variation giving rise to social organization such as leadership. A specific form of leadership can emerge under unstable environmental conditions, when some individuals become better suited than others to initiate action and influence the behavior of their peers. Unlike fixed dominance hierarchies, emergent leadership can rapidly adjust to changing environmental conditions, thereby reshaping group organization. Because it does not require the maintenance of stable hierarchies, this form of leadership can arise even in species that do not have the cognitive capabilities to sustain complex social structures. In this work, we investigate the combined effects of individual heterogeneity and emergent leadership on the evolution of cooperation using an evolutionary game-theoretic model in which individuals may assume the roles of leaders or followers according to their strength, representing individual differences in suitability to prevailing environmental conditions. We examine different levels of population heterogeneity together with increasingly complex strategy sets requiring progressively greater informational requirements, allowing individuals to condition cooperation on their own strength, leadership role, or both. Our results show that the interplay between leadership and heterogeneity promotes the evolution of cooperation, particularly when only a small fraction of individuals act as leaders. Under these circumstances, cooperation evolves even when individuals employ the simplest possible strategies. Under harsher ecological conditions, cooperation can be sustained by more sophisticated strategies, specifically by conditional strategies that prescribe cooperation when individuals are strong or leading and defect when acting independently.

**Author summary:** In this study, we propose that emergent leadership mediated by individual diversity can boost the evolution of cooperation in animal groups. Building on growing evidence on the heterogeneity of animal capabilities and personalities, we focus on the fleeting leadership that emerges in animal groups when facing rapidly changing environmental conditions. We suggest that this type of leadership that emerges from individual differences in strength—a generic quality encompassing those characteristics that make an individual more fit to lead in a given situation—does not require complex cognitive capabilities from the animals and represents a valid alternative to more demanding strategies proposed in the past to explain the evolution of cooperation. Using an evolutionary game theory model, we show that if a population includes a few strong players, these can become influential leaders and guide the actions of their peers to achieve cooperation. Although the naive strategy of always cooperating is sufficient for cooperation to evolve, the introduction of more complex strategies leads players to cooperate only when they are more likely to be recognized as influential leaders. These strategies are more effective in promoting cooperation under unfavorable ecological conditions and are also more robust against exploitation by defectors.

## Introduction

Although empirical evidence has shown that kin selection can explain cooperation in bacteria [1–3], eusocial insects [4], and some larger-brained social species, challenges arise when studying cooperation between unrelated individuals [5]. Many mechanisms have been suggested to explain the evolution of cooperation among non-kin, but several of the most influential ones require non-trivial cognitive abilities. For instance, for reciprocity to be possible, animals should be capable of mate recognition and long-term memory, especially when interactions occur within large groups [6]. Other studies have suggested that signaling techniques can support the evolution of cooperation [7, 8], although this shifts the explanatory challenge to understanding how reliable communication systems themselves evolve.

One aspect that has historically received less emphasis in studies of animal cooperation is the role of population diversity.. Diversity has been shown to be widespread within animal species and phenotypic variation is selectively maintained in populations [9]. Among the behavioral traits commonly reported to vary within populations are those associated with leadership (e.g., boldness or aggressiveness).

Leadership is one of the most widely debated topics when studying animal collective behavior [10–12]. It plays a key role in decision making, i.e., leading to group movements [13–15]. However, not many studies explicitly include leadership as a mechanism able to influence the emergence of cooperation, and, when they do, the focus is on rigid social hierarchies that are only displayed by a narrow range of animal species [16]. Empirical evidence suggests that, in many species of animals, leadership is not enforced by a rigid hierarchy, but emerges dynamically, as the environmental conditions require, from interactions among phenotypically and behaviorally heterogeneous members of the group [17].

Leadership in animal groups is usually classified either as “unshared” or “distributed” [18]. In the former, an individual leads the group by imposing their will in a despotic way or by virtue of their role in the social hierarchy. The latter, instead, entails an organization in which any member has the possibility of assuming the leading role [19, 20]. Sometimes, the emergence of the one or the other category depends on the characteristics that make an individual a leader, including its phenotype, the group composition, and the environmental conditions. One strong animal might naturally become the only leader within a group of weaker members but would share the leadership within a group formed by equally strong individuals. The distributed leadership is particularly favorable when contingencies faced by animal groups vary frequently. Here, rigid social hierarchies may hinder the surviving capability of the group: since group leaders are generally individuals particularly sensitive to environmental stressors and suitable for certain specific conditions, environmental change can significantly affect the behavior of the groups and the effectiveness of their leadership hierarchies [9]. Instead, certain environmental contingencies can make some individuals especially adapt to assume leading positions in the group. These individuals can, in turn, influence the behavior of their group mates more or less effectively. This fleeting hierarchical order is inherently linked to the situation the group is dealing with, and as this changes, it will change accordingly, with other animals assuming the leading role.

There exist several ways in which transient leadership can arise in combination with heterogeneity. For example, individuals with higher locomotor and metabolic capacities and those with less social and more goal-oriented behavior can act as leaders that encourage group movements [9, 21]. In some species, leading is associated with changing energy requirements [22, 23]: unfed caterpillars are more likely to initiate foraging bouts [24], hungry individuals occupy front positions in fish shoals until they regain their nutritional balance [25]. Some animals may assume the leader role only when paired with less bold individuals, and boldness is highly dependent on past experience [26]. Not only do bolder individuals tend to act as leaders, but shyer individuals tend to elicit greater leadership tendencies in their bold partners [27]. Physiological differences might promote some individuals to the leader role in some species: in pigeons, leadership hierarchies are the consequence of differences in birds’ speed [28]; leading and following behavior are modulated by fish size in shoals of golden shiners [29]; and schooling fish form temporal leadership interactions based on the relative speed between neighbors [30]. In some species, older members lead the group when more knowledge is advantageous, such as in uncertain situations [31, 32].

Furthermore, individual behavioral differences have been shown for many different behaviors, including those often associated with leadership such as boldness and aggression, and in a wide range of species, from microbes to humans [33, 34].

Importantly, leadership roles are not always associated with dominance hierarchies but can reflect a distinct form of social organization shaped by cooperative demands rather than preexisting power dynamics [35].

The fact that cooperation is achieved by animals with very different cognitive abilities, even minimal ones [5, 36], calls for the study of mechanisms that sustain cooperation that do not require any individual sophisticated ability. As shown above, heterogeneity and emergent leadership are ubiquitous properties across the animal kingdom that arise even among the simplest organisms, as a result of their diversity. A distributed leadership emerging from group heterogeneity does not require social relations to be maintained and enforced, and thus removes the need for complex cognitive capacities that are essential to sustain social intelligence. For these reasons, heterogeneity and leadership might offer critical insights into the evolution of cooperation, even in animals with limited cognition. Indeed, there is evidence that individuals of many species, including parakeets [37], vervet monkeys [38], ravens [39], fish [40], and baboons [41], establish their role in the social order of the group simply by interacting with others and observing others interacting. Moreover, dominant and subordinate behaviors do not always require physical interactions, but can be based on cues about another individual’s competitive ability, called status signals [42, 43].

Through these mechanisms, individuals can assess their role in the group and use this knowledge to inform their behavior.

Here, we propose an evolutionary game model to study the conditions for the evolution of cooperation in heterogeneous groups of agents when shared resources and collective benefits are at stake. In this model, players engage in public goods games in which they have to make decisions about collective actions by which personal incentives may conflict with the common good. Public goods games are widely used across different disciplines since they constitute a flexible and powerful framework to study the foundations behind cooperation and collective behavior [44, 45]. Taking into account different degrees of heterogeneity and sets of strategies relying on increasingly demanding cognitive skills such as awareness of social strength and role in the group, our model offers insights into the intertwined dynamics between heterogeneity and distributed leadership and how these two influence the evolution of cooperation under many different conditions. We demonstrate that in a heterogeneous population, strong individuals can become influential leaders and promote the evolution of cooperation even under unfavorable conditions. As conditions worsen, unconditional cooperation is no longer sufficient, and more complex strategies are required to sustain it. Among these, we find that the most successful are those that make individuals cooperate only when they are influential leaders, thereby avoiding exploitation by defectors while securing the cooperation of their followers.

### Model

In order to study the emergence and evolution of cooperation in leadership contexts, we take a game-theoretical approach. Two time scales are considered: a faster one, where individuals interact according to game rules and individual strategies within a population, and a slower one, where evolutionary dynamics take place.

### Game Definition

A commonly-used game to represent cooperation in a group context is the public goods game [46]. Here, we consider groups of *N* players randomly selected from the population, who interact choosing either to cooperate or to defect. Cooperating players invest a fixed amount *c* in a common pool, while defecting players do not incur any cost. At the end of each round, all players receive the same benefit, obtained by multiplying the total amount invested by cooperators by a factor *r* and dividing it by the total number of players. This multiplication factor *r* represents the efficiency with which individual cooperative efforts are transformed into common benefits, and thus serves as an indicator of how favorable ecological conditions are. As a consequence, each player receives a benefit 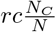, which is proportional to the number of cooperators in its group *N*_*C*_. Considering that each individual *i* cooperates with a probability *P*_*i*_(*C*), the average payoff obtained upon interaction in a given group is

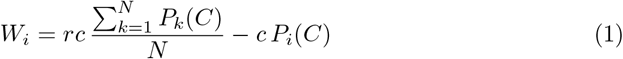

Each player has the opportunity to become a leader in its group. Leaders play according to their own intrinsic strategy, while non-leaders can either stick to their own strategy or follow a leader by replicating its action. Both becoming a leader and following a leader depend on the *strength* of the individuals within the group. Fig 1 schematically represents how the leadership mechanism works. Every individual in the population has a probability *p*_*s*_ of being *strong* (*s*) and a probability 1*−p*_*s*_ of being *weak* (*w*). This probability can also be interpreted as the fraction of individuals that on average are strong in the population. Each strength level *σ*_*i*_ *∈{s, w}* that individual *I* exhibits is linked to a strength value 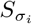, defined by a Fermi function as follows:

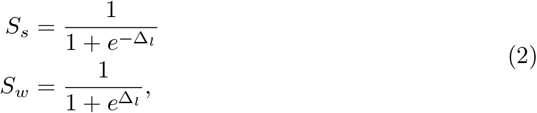

where Δ_*l*_ determines how different the strong and weak individuals are. Hence, Δ_*l*_ represents the heterogeneity of the population in terms of strength.

**Fig 1.**
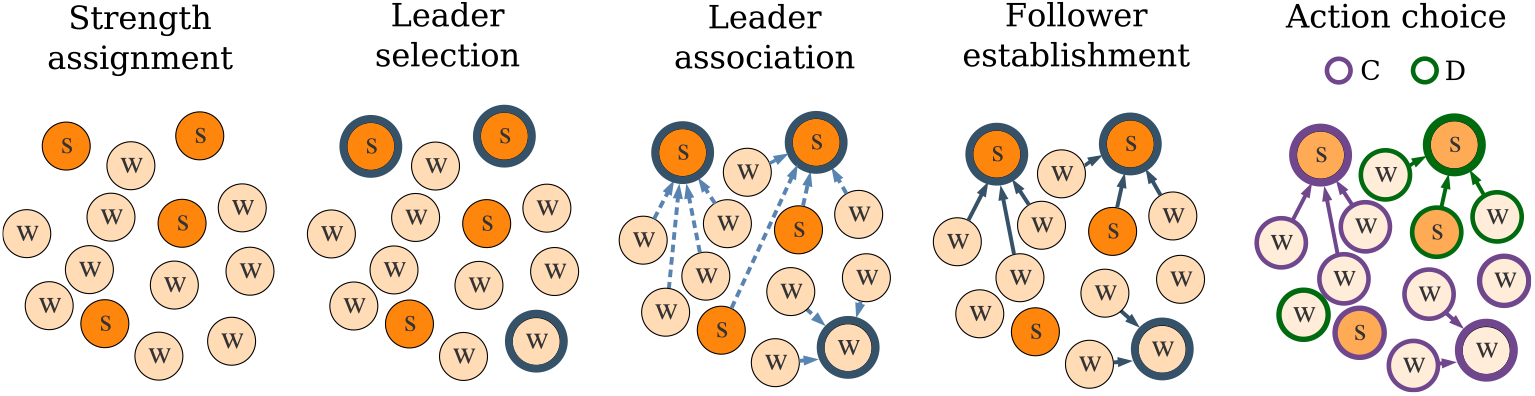
Leadership mechanism. Conceptual illustration of the leadership mechanism operating within a group.

We assume that the probability of a player *i* with a strength level *σ*_*i*_ becoming a leader is equivalent to the strength value:

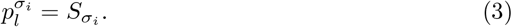

Therefore, as Δ_*l*_ increases, strong and weak players become, respectively, more and less likely to act as leaders. Note that in the rare cases where no leaders are stepping up or all individuals become leaders, the leadership mechanism is not taking place, and all individuals play their own strategy.

Once leaders are chosen among the individuals that constitute the group, each non-leader individual *j* is assigned to a leader *i* proportionally to the strength level 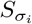, with probability 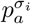:

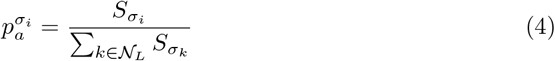

where *N*_*L*_ is the set of leaders in the group. Note that the higher the leader strength, the higher the number of potential followers. Each non-leader *j* further determines whether or not to finally follow its assigned leader *i* as a function of their strengths. Since there only exist two strength levels, we establish four probabilities 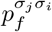of a non-leader with strength level *σ*_*j*_ to follow a leader with strength level *σ*_*i*_:

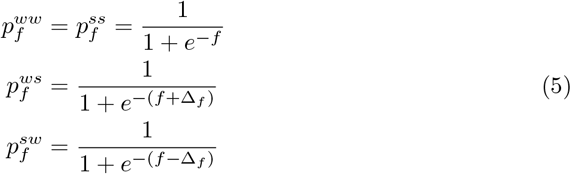

These probabilities are expressed as Fermi functions, where *f* represents the general tendency (intensity) of the population to follow the leaders, and Δ_*f*_*≥*0 modifies this tendency when the leader and its potential follower have different levels of strength. If Δ_*f*_ = 0, there is no heterogeneity of following behavior in the population: once a leader is assigned to a non-leader, the probability of the latter following the former does not depend on their strength. When increasing the value of Δ_*f*_, weak players become more likely to follow strong leaders and strong players become less likely to follow a weak leader. Since both Δ_*l*_ and Δ_*f*_ represent population heterogeneity, it is reasonable that they are related to each other. Hence, we focus first on the case where Δ_*l*_ = Δ_*f*_ =: Δ. Joining Eqs 3, 4 and 5, the probability that a player *j* with strength level *σ*_*j*_ follows a player *i* with strength level *σ*_*i*_ can be synthesized as:

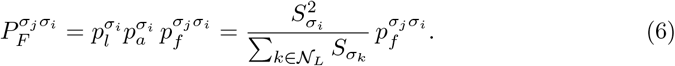

We also study a set of scenarios in which, among all members of the group, one and only one individual emerges as a leader [47]. In a single-leader scenario, the probability that an individual *i* with a strength level *σ*_*i*_ becomes the sole leader is proportional to its strength value, but normalized to the sum of the strengths of all players in the group.

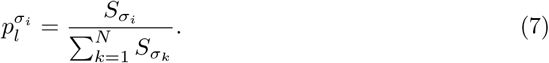

Since there exists only one leader in a group, the rest of individuals are non-leaders and assigned to the former, such as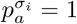. Therefore, Eq 6 becomes in the single-leader scenario:

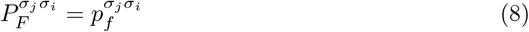

### Strategies

We study four scenarios in which different sets of strategies are in place at increasing levels of complexity. Each strategy is defined by a set of bits that indicate the probability of cooperation in given circumstances. Although we write these probabilities as 1 (cooperation, **C**) or 0 (defection, **D**) for simplicity, they are implemented as 1*− ϵ* and *ϵ*, respectively, where *ϵ* is the small probability of acting contrary to the intended action. Given that our results are not significantly affected by the specific value of this error when it remains low, we assume *ϵ* = 0.01 from now on.

The simplest set of strategies is defined with only one bit for the action of the player *a*, and includes always cooperate **AllC** [1] and always defect **AllD** [0]. This, hereafter referred to as the **B** scenario, serves as a baseline for analyzing how increased complexity affects the emergence of cooperation within a leadership context.

In the next level of complexity, individuals decide their *free* actions conditional on their awareness about their strength (weak or strong) or their role (leaders or not). This gives two possible sets of strategies of two bits each. On the one hand, strategies conditional on the individual strength level—from now on referred to as the scenario **S**—are defined as [*a*_*w*_, *a*_*s*_] and include, together with **AllD** [00] and **AllC** [11], a strategy that makes individuals cooperate when they are weak and defect when they are strong (**WCSD** [10]), and the opposite strategy that makes individuals cooperate when they are strong and defect otherwise (**WDSC** [01]). On the other hand, strategies conditional on the individual role—hereinafter called the **L** scenario—are defined as [*a*_*nL*_, *a*_*L*_], and include **AllD** [00], **AllC** [11], **NCLD** [10], which makes individuals cooperate when they are not leading and otherwise defects, and **NDLC** [01], which makes individuals cooperate only when leading.

Finally, the two sets of two-bit strategies are combined in a single set of four-bit strategies—referred to as the **S+L** scenario—defined as [*a*_*w,nL*_, *a*_*s,nL*_, *a*_*w,L*_, *a*_*s,L*_]. Each bit represents the action taken by individuals when leading or not while being either weak or strong. For example, *a*_*w,L*_ is the probability that a weak leader cooperates.

Previously defined strategies have their equivalents in this set: **AllC** [1111], **AllD** [0000], **WCSD** [1010], **WDSC** [0101], **NCLD** [1100] and **NDLC** [0011]. Due to their importance, we identify the strategy that makes only strong leaders cooperate as **SLC** [0001] and the group of strategies that makes any leader cooperate as **LC** [0*11], where * includes both cooperation and defection.

### Evolutionary dynamics

Evolution is modeled using a stochastic birth-death process in a well-mixed finite population of size *Z* [48]. Individuals randomly assemble into groups of *N* players and repeatedly engage in interactions governed by the game described above. The average payoff of each individual, calculated in all possible group configurations, serves as a measure of fitness. Reproduction favors fitter individuals, with the transmission of a strategy following a Fermi function controlled by the intensity of selection *β* [49]. This ensures that offspring inherit behavioral rules for both strong and weak, and for leader and non-leader rounds, with the probability of being strong or weak remaining fixed throughout evolution. The calculation of payoffs and fixation probabilities is further detailed in Materials and methods and a summary of the main parameters of the model can be found in Table 1.

**Table 1.** Main parameters of the standard model.

| Parameter | Description | Value |
| --- | --- | --- |
| $Z$ | Population size | 100 |
| $N$ | Group size | 9 |
| $c$ | Cost of cooperation | 1 |
| $r$ | Benefit multiplication factor | $[1, 10]$ |
| $p_s$ | Probability of players to be strong | $[0, 1]$ |
| $\Delta_l$ | Difference in strength value between strong and weak individuals | $[0, 5]$ |
| $f$ | Tendency of the population to follow a leader | 0 |
| $\Delta_f$ | Difference in tendency to follow a leader between strong and weak individuals | $[0, 5]$ |
| $\epsilon$ | Action error probability | 0.01 |
| $\beta$ | Intensity of selection in evolutionary dynamics | 1 |

## Results

### Leadership and self-awareness promote cooperation in heterogeneous populations

Following the evolutionary dynamics described in detail in Materials and methods, the stationary distribution of strategies has been computed in finite populations under different scenarios. Fig 2 shows the average level of cooperation obtained from these stationary distributions as a function of *p*_*S*_ and different values of *r* and Δ. While in standard public goods game defection prevails when *r < N* [46], the introduction of the leadership mechanism promotes the evolution of cooperation even for low values of *r*. This is evident in the baseline scenario **B** when ecological conditions are more favorable (intermediate or high *r*), especially for high values of Δ, which implies large differences in strength between strong and weak players. The awareness of the own strength and leadership role (**S, L** and **S+L** scenarios) further improves the levels of cooperation even for less favorable conditions (e.g., *r* = 3 in Fig 2) and for low values of Δ.

**Fig 2.**
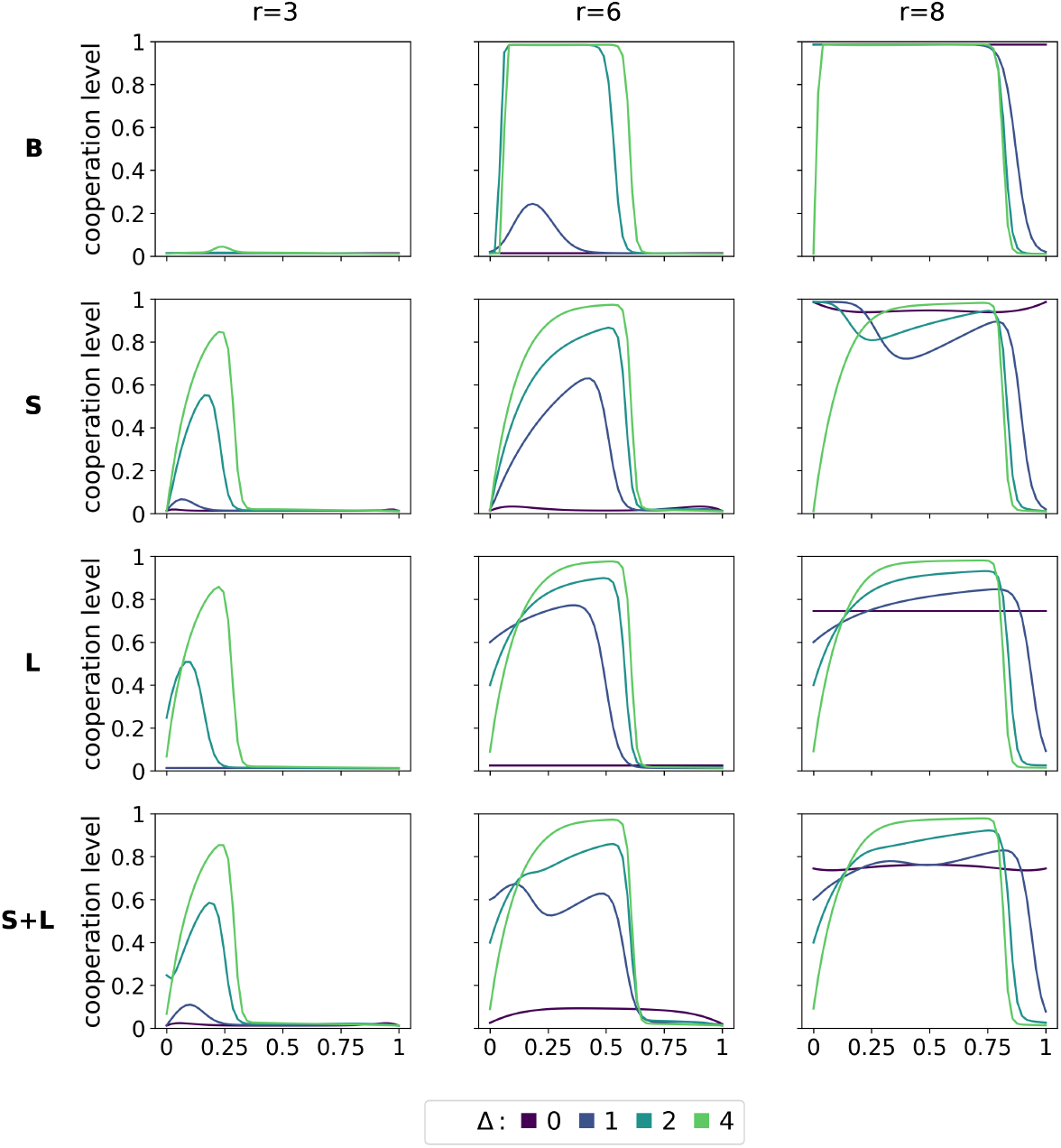
Leadership promotes the evolution of cooperation. Cooperation levels as a function of *p*_*S*_ and Δ for three representative values of the reward factor *r* across the different strategy scenarios. Fixed model parameters are *Z* = 100, *N* = 9, *f* = 0, *ϵ* = 0.01, and *β* = 1.

The high levels of cooperation observed when Δ is large indicate that the leadership mechanism works effectively. Under this regime, the level of cooperation increases with the probability of being strong, *p*_*S*_, until it reaches a maximum. Beyond this point, cooperation collapses for all values of *p*_*S*_, exhibiting a behavior reminiscent of a phase transition between the cooperation and defection regimes. This phase transition implies that a very small change in the probability of being strong can determine whether a population remains highly cooperative or becomes fully defective. The threshold value of *p*_*S*_ was analytically approximated in Materials and methods for the high-heterogeneity regime, which yields the condition for the maintenance of cooperation:

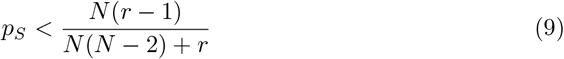

This analytical expression is well supported by the numerical results. For instance, when *N* = 9 and *r* = 6, Eq 9 predicts that cooperation is promoted for *p*_*S*_ *<* 0.65, in agreement with the transition observed in all four scenarios shown in Fig. 2. As further predicted by Eq 9, Fig. 2 and Fig. S1 show that the threshold value of *p*_*S*_, and consequently the overall level of cooperation, increases with *r* and decreases with group size.

Low values of Δ imply a less effective leadership mechanism, either because not all strong individuals become leaders while some weak individuals do (low Δ_*l*_), or because following behavior is more erratic and less strongly associated with individual strength (low Δ_*f*_ ). In Materials and methods, we analytically derive the conditions under which cooperation can be maintained when Δ = 0, showing that cooperation is guaranteed only under favorable ecological conditions. Fig. 2 shows that under this regime the level of cooperation is reduced, except for very high or very low values of *p*_*S*_ when *r* is high. Only in these limited situations, some cooperation is guaranteed if leaders and the decision to follow them are taken randomly.

In the extreme situation of a uniformly weak population (*p*_*S*_ = 0), a lack of leaders is expected, and even when some individuals occasionally assume this role, their ability to influence others is limited. On the other hand, when all individuals are strong (*p*_*S*_ = 1), most individuals become leaders, leaving very few, or none at all, available to act as followers. In both cases, the leadership mechanism becomes ineffective and the game effectively reduces to a standard public goods game, where defection easily takes over the population.

To understand what makes leaders effective, we also studied the case in which Δ_*l*_ and Δ_*f*_ are not equal. Under these conditions, we found that the former has a stronger impact in shaping cooperation levels than the latter (see Fig. S2 and Fig. S3, respectively).

The presence of multiple leaders in a group—especially with high values of *p*_*S*_—leads to diluted following and consequently to a reduction in the chances of cooperation. In contrast, when there is one and only one leader, as studied in the single-leader scenario (see Fig S4), such conflicts do not emerge, leading to a boost in cooperation. Although this effect is already present in the **B** scenario, it is more evident in the **L** and **S+L** scenarios where individuals are aware of their role as leader. This awareness, combined with the clearer role of the only leader that does not have to compete with others for the attention of followers, guarantees high levels of cooperation even when the leader is not that strong (low Δ). Moreover, the single leader scenario leads to increased cooperation in the extreme cases where everyone is weak (*p*_*S*_ = 0), always ensuring the presence of a leader, and when everyone is strong (*p*_*S*_ = 1), removing the competition between multiple leaders. For this reason, the **L** and **S+L** scenarios do not show a phase transition between the cooperation and defection regimes. Interestingly, the level of cooperation reaches its maximum when the probability of being strong (*p*_*S*_) is relatively low (around one quarter of the population for the combination of parameters displayed), especially compared to the multiple leader case, and decreases smoothly when *p*_*S*_ increases, maintaining a relatively high level of cooperation for all combinations of parameters studied. In single-leader scenarios, it is more advantageous that there exist a few strong individuals, just enough to avoid groups without a strong leader, but not many to ensure that most individuals follow the strong leader. The scenario **S** benefits less from the simpler leadership mechanism of the single leader case, because strength awareness alone does not grant those individuals that become leaders the ability to fully exploit their leadership role.

The results presented above are robust with respect to several parameter variations. As already mentioned, the model was tested against variations in group size *N* (see Fig S1). Increasing *N* reduces the *p*_*S*_ threshold under which cooperation dominates in multiple-leader scenarios according to Eq 9. The general level of cooperation is also reduced when Δ is low for multiple-leader and **S** single-leader scenarios. In Fig S5, we study scenarios where individuals have a higher probability *ϵ* of making action errors, showing that increased errors are associated with a decrease in the level of cooperation. However, in most cases, no significant differences are noticeable on a qualitative level. Only in single-leader scenarios in which individuals cannot make their decisions based on their leadership role (i.e., **B** and **S** scenarios), cooperation suffers a severe reduction, since mistakes of a sole leader can spread quickly. Both these robustness analyses suggest that when leadership and general role behavior are more clear, i.e., single-leader scenarios with individuals aware of their leadership roles and acting accordingly (**L** and **S+L** scenarios), systems are more resilient to higher error rates and can foster cooperation in larger groups.

### Role of leadership strategies in promoting cooperation

In order to better understand the dynamics underlying the results previously shown and the role of the different behaviors in play, we analyze the stationary distributions for different scenarios and the dynamics that give rise to them. Fig 3 shows the probabilities in the stationary distribution of the main strategies that emerge in the different multiple-leader scenarios and Fig 4 displays the invasion graphs for some representative cases indicated by the yellow dots in Fig 3 (see Figs S6–S7 for the stationary distribution of all strategies, and Figs S8–S9 for other invasion graphs). The phase transition between cooperation and complete defection observed in Fig 2 and analytically approximated in Eq 9 is clearly reflected. As observed in Fig 3, as Δ decreases, the phase transition between cooperative strategies (**AllC, WDSC**, or **NDLC**) and **AllD** occurs for lower values of *p*_*S*_.

**Fig 3.**
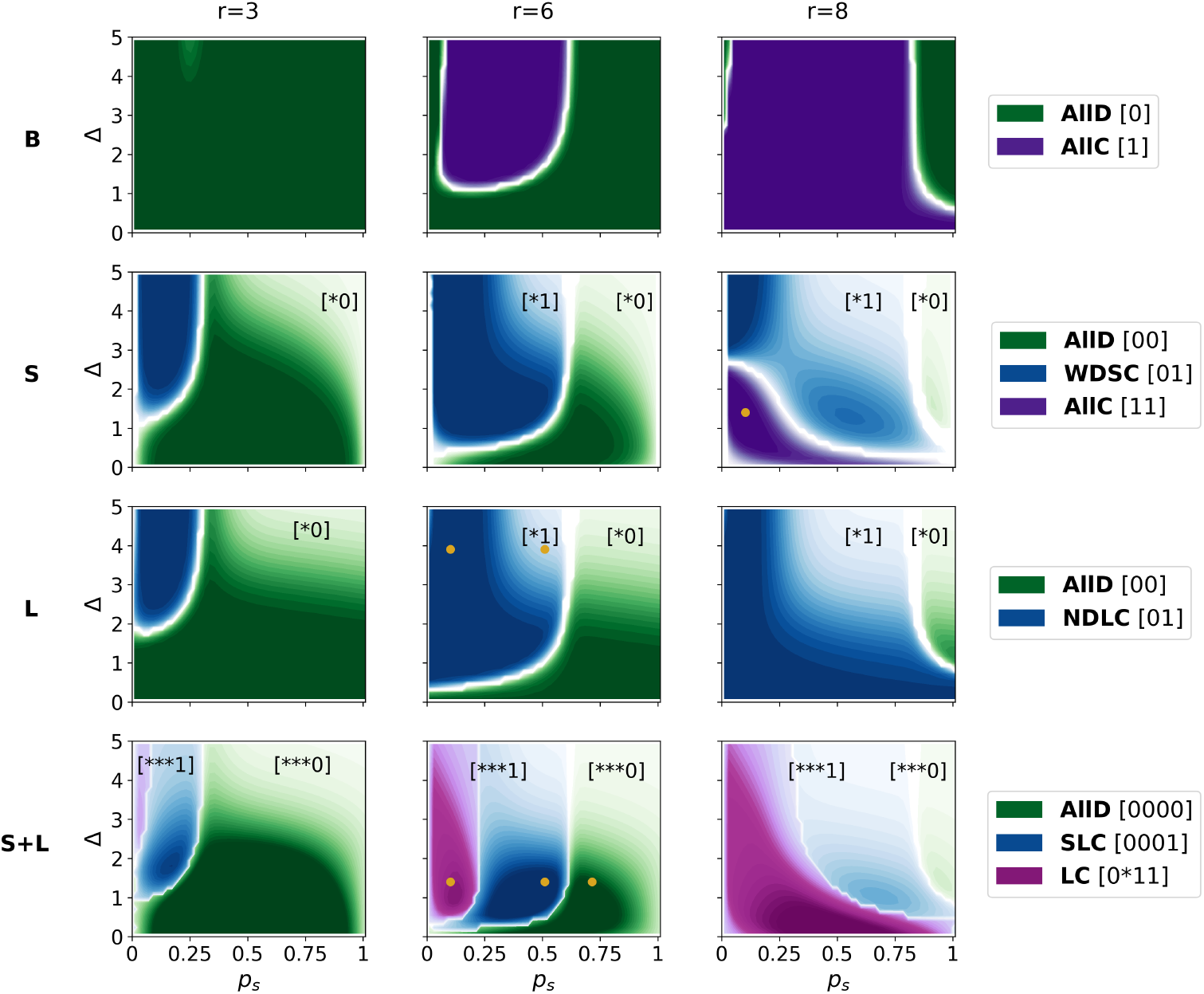
Leadership behavior determines the emergence of cooperation and defection. Stationary distributions of the dominant strategies for three representative values of the reward factor *r* across the parameter space defined by *p*_*S*_ and Δ, for each strategy scenario. For clarity, only probabilities exceeding 0.5 are shown in the **B, S**, and **L** scenarios, whereas all probabilities greater than 0.125 are shown in the **S+L** scenario. Fixed model parameters are *Z* = 100, *N* = 9, *f* = 0, *ϵ* = 0.01, and *β* = 1. Yellow dots indicate the parameter combinations used to generate the invasion plots shown in Fig. 4.

**Fig 4.**
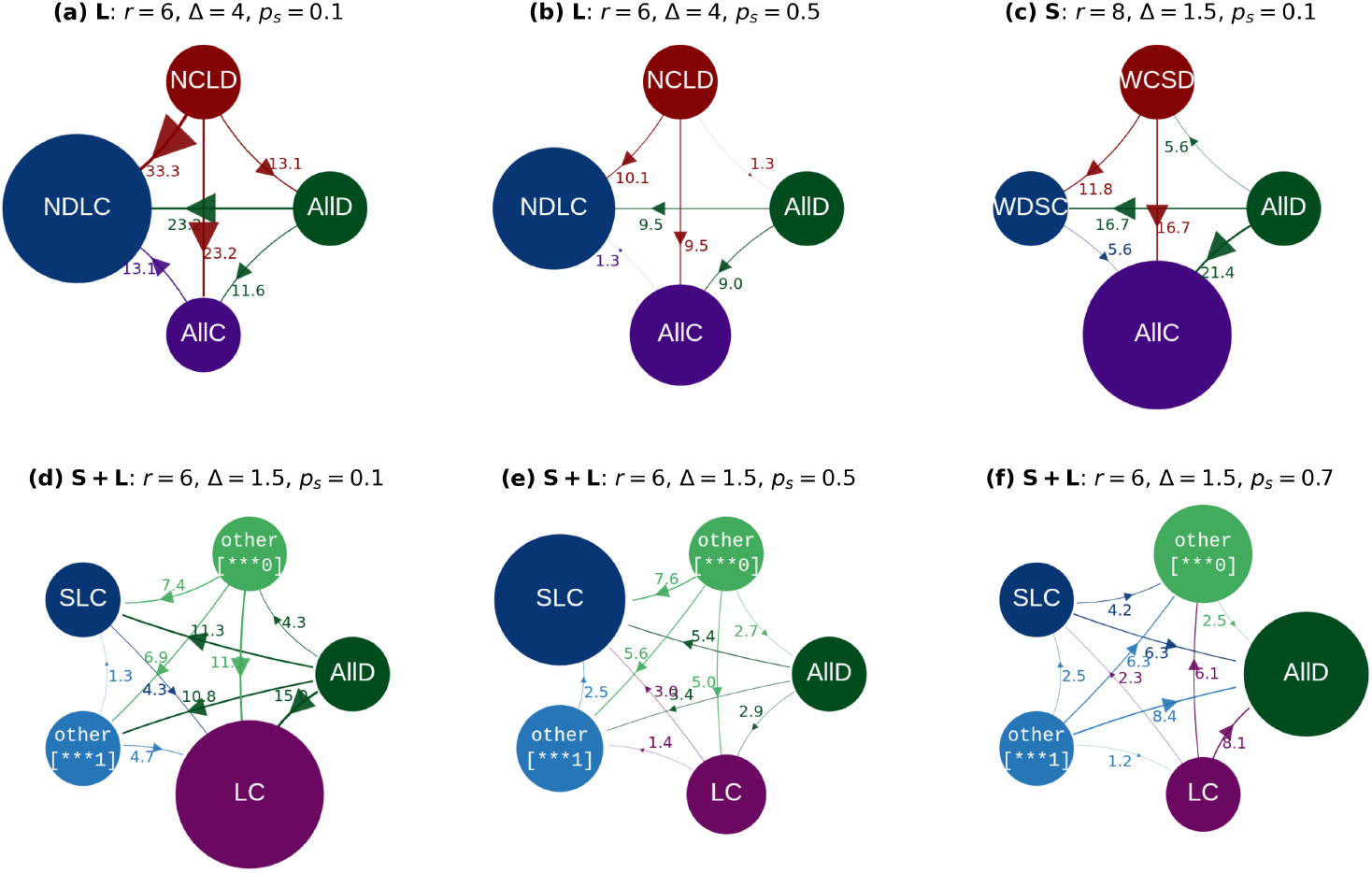
Invasion graphs of the main strategies. Invasion graphs are obtained for different strategy scenarios and representative values of the reward factor *r*, corresponding to selected regions of the parameter space defined by *p*_*S*_ and Δ marked in Fig. 3. Node sizes are proportional to the probabilities in the stationary distribution. Arrow widths and their associated numerical values represent transition probabilities between strategy groups, normalized by neutral drift and, when applicable, by the stationary distribution of the receiving group. Fixed model parameters are *Z* = 100, *N* = 9, *f* = 0, *ϵ* = 0.01, and *β* = 1.

The most important insight from Fig 3 is that awareness of one’s own strength and/or leadership role is crucial for keeping defectors at bay under unfavorable conditions (i.e., low values of *r*). Indeed, when individuals lack this capability, as in the **B** scenario, **AllD** dominates **AllC** across a broader range of parameter values. In the other scenarios, on the contrary, the evolution of cooperation depends on how strong and leading individuals behave, as reflected in the dynamics observed before the critical thresholds separating cooperation from defection. Across all these scenarios, conditional cooperators outperform unconditional cooperators. Strategies such as **WDSC, NDLC, SLC**, and **LC**, in which individuals cooperate only when they know that they are leaders or are likely to become leaders, are stronger than unconditional cooperation (**AllC**). These strategies resist invasion by defectors by maintaining high levels of cooperation when acting as leaders while defecting when not in leadership positions, thereby exploiting more naive cooperators (see Figs. 4 and S8–S9). Indeed, when strong leaders are rare (low *p*_*S*_ and high Δ), many weak, non-leading individuals defect, reducing overall cooperation and making **AllC** vulnerable to invasions (Fig 4a). In contrast, when groups are more likely to include strong influential leaders (high *p*_*S*_ and high Δ), individuals that cooperate when they know that they are leaders or are likely to become ones (**WDSC, NDLC** and **AllC**) assume leadership roles more often, thereby fostering higher levels of cooperation (Fig. 4b).

In parameter regimes where strength and leadership are weakly correlated (low Δ) and the population contains too few or too many strong individuals (low or high *p*_*S*_), cooperation evolves only under highly favorable conditions (high *r*). The threshold value of *r* required for cooperators to invade defectors was derived in Eqs. 31 and 32 in Materials and methods. Interestingly, when strong individuals are scarce (low *p*_*S*_) in the **S** scenario, conditional cooperators, such as **WDSC**, only spread defection. As a result, under favorable conditions (high *r*), unconditional cooperation (**AllC**) becomes competitive again, as shown in Figs 3 and 4c. By contrast, in the **L** scenario, **NDLC** continues to outperform full cooperation, as individuals adopting this strategy are capable of taking advantage of their leadership position regardless of whether they are weak or strong

The results of the **S+L** scenario largely mirror those observed in the **S** and **L** scenarios (Fig 3). In particular, **SLC** and **LC** play roles analogous to those of **WDSC** and **NDLC**, respectively. Individuals adopting **SLC** and **LC** strategies cooperate when they know they are being followed (i.e., when they are strong leaders) and defect when they are weak non-leaders. As in the other scenarios, when *p*_*S*_ and Δ fall and the number of weak individuals becomes significant, more of them will also become leaders. Under these conditions, cooperating only when acting as a strong leader (**SLC**) becomes overly exploitative and yields lower fitness, allowing more cooperative strategies, such as **LC**, to dominate (Fig 4d). By contrast, increasing *p*_*S*_ and consequently the likelihood of strong leaders in the group allows **SLC** to dominate the population, maintaining high levels of cooperation until the phase transition to full defection (Fig 4f).

In the single-leader case, Fig S11 shows that cooperative strategies, especially **AllC**, emerge across a broader range of parameter values in the **B** and **S** scenarios than in their multiple-leader counterparts. Due to the undisputed role of the sole leader, unconditional cooperators (**AllC**) are less vulnerable to exploitation. By contrast, the results for the **L** and **S+L** scenarios explain the absence of a phase transition in the cooperation level. In these cases, the stationary distribution is dominated by the strategies that most effectively exploit the single-leader mechanism, namely **NDLC** and **LC**. Although the multiple-leader scenario can reach higher peaks of cooperation, it is also more prone to collapsing into full defection if the proportion of strong individuals is too high.

## Discussion

Heterogeneity and leadership can be an effective evolutionary mechanism underlying cooperation in heterogeneous populations. Leadership, even in its simplest forms, is able to promote cooperation: when ecological conditions are highly favorable but not sufficiently to ensure cooperation in the absence of additional mechanisms, the random emergence of leaders and their followers already ensure a certain level of cooperation. However, as conditions become more severe, leadership must be associated with heterogeneity-driven expression of different roles, and the ability to act conditionally to the own role plays a decisive role. Indeed, populations in which individuals are aware of their own strength and/or leadership roles are able to maintain cooperation even under the most unfavorable conditions.

The results show that the behavior individuals adopt when they recognize themselves as leaders is the key factor determining whether a population manages to maintain cooperation or, on the contrary, is drawn towards defection. Particularly, individuals who invest in the public good when they become influential leaders (or have high chances to become one) and defect when they are by their own—not following or being followed—are key to sustain cooperation. They can be seen as a lesser evil capable of keeping defection at bay when ecological conditions are not favorable or heterogeneity is not high, since they are more resistant to being invaded by defectors. This evokes a similar conclusion from works on cooperative agreements [50] and signaling [8], which shows that cooperation needs individuals willing to contribute exclusively when necessary.

Our analysis also shows that there exists an interval in the fraction of strong individuals in the population for which cooperation can emerge. In particular, when there is no limitation in the number of leaders allowed, cooperation increases with the number of potential leaders until a critical threshold—which we deduced analytically for high heterogeneity—where a phase transition occurs and full defection takes over. These findings offer a complementary explanation for the frequent occurrence in nature of groups in which only a few individuals happen to lead, which is typically attributed to the elevated evolutionary costs associated with developing stronger capabilities.

Therefore, evolution favors the existence of a minority of strong leaders within a majority of weaker potential followers, thereby promoting cooperation at lower cost and conflict.

We also studied systems in which only one leader is allowed. This constraint may arise from cultural or biological factors (e.g., genetically determined roles), as well as from decision-making dynamics (e.g., the first individual to act becomes the potential leader). These systems are much more effective at coordinating groups toward cooperation under high uncertainty—such as low heterogeneity or high error rates—or when ecological conditions are unfavorable. Furthermore, when individuals are aware of their leading roles, single-leader systems manage to preserve relatively high general cooperation besides variations in surrounding features, since they grant certainty and avoid fragmentation. However, when conditions are favorable, they are not able to reach cooperation levels as high as those of multiple-leader systems.

The model developed in this work is meant to study leadership as a mechanism to promote cooperation from a fundamental point of view, and it is not aimed at covering the intrinsic particularities of the countless systems where it appears. We have assumed that the chances of leading are tied to the strength of the individual. However, if individuals obtain leadership positions through other means rather than their own competence, as sometimes happens in more complex groups such as human societies (e.g., political populism, social media influencers, etc.), the number of weak leaders may become significant and leadership mechanisms can lose their effect on cooperation, allowing defection to take over (e.g., distrust of authorities and political systems). The model does not take into account either that leaders could obtain different benefits and costs as a function of their success in recruiting followers and/or fulfilling global benefits for the group, which would imply higher individual cognitive capacities.

With regard to human groups, the results of this study may also help inform management policies. Individuals who are usually perceived as defectors may in fact behave in this way because they are following a **NDLC**-like strategy, possibly without being conscientious of it. Their behavior could then change when they are entrusted with leadership responsibilities. This approach offers an alternative or complement to direct punishment, ostracism, or reward, which are often costly and inefficient, particularly in conflict-prone contexts (e.g., schools, companies, and rundown neighborhoods). On the other hand, our results suggest that societies which display uncertainty on individuals’ power roles, where competence does not clearly translate into leadership positions, or simply where economic conditions are not favorable, can particularly benefit from making individuals aware of their potential and capabilities, ultimately contributing to the empowerment of democracy.

## Materials and methods

### Analytical model development

Here we develop in more detail how payoffs and stationary distributions can be calculated for the four-bit strategy set, which constitutes the most general scenario our model is covering. The analysis of the other sets of strategies can be developed as simplifications of this one.

To simplify the evolutionary analysis, the small mutation approximation is adopted, where a single mutant either fixates or disappears after invading a uniform population. This reduces the evolutionary dynamics to transitions between monomorphic states without the possibility of mixed states, which could lead to a much more complex ecosystem of strategies [51, 52]. Under this approximation, only two strategies are present in the population at most. Following Eq 1, the payoff that an individual playing strategy A obtains when in a group formed by *N*_*A*_ individuals playing this strategy and *N − N*_*A*_ playing another strategy B can be computed as

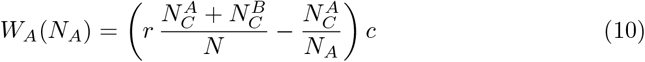

where 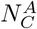and 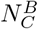are the number of cooperators playing strategies A and B, respectively, and can be calculated as

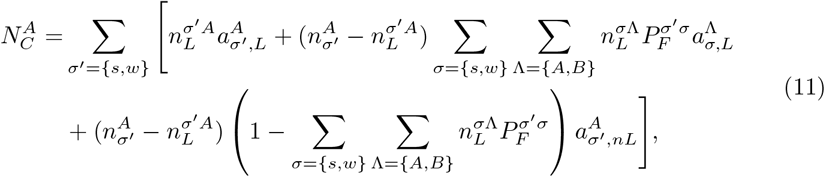

where 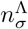 represents the number of individuals with strength level *σ* = *{s, w}* who play strategy Λ= *{A, B}*, of whom 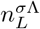 are leaders. Note that 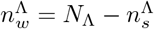. In the previous equation, the first term counts for individuals with strategy A becoming leaders; the second term counts for they following leaders (using Eqs 6 or 8), and the last term counts for they neither becoming leaders nor following any other individual. Note that in order to calculate the payoff for strategy B, A and B must be switched in the previous equations.

Assuming that there exist *Z*_*A*_ individuals playing strategy A and *Z− Z*_*B*_ playing strategy B in the population, the average payoff of each strategy Λ= *{A, B}* over all group combinations is computed as

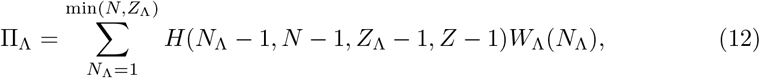

where the hypergeometric distribution can be expressed as

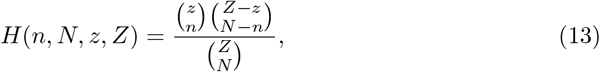

and *W*_Λ_(*N*_Λ_) represents the payoff that individuals playing strategy Λobtain when in a group formed by *N*_Λ_playing this strategy and *N−N*_Λ_individuals playing the other strategy. Counting for all possible combinations of individual strength levels in the group, this payoff is calculated as:

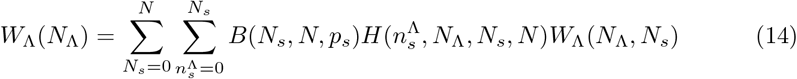

where *N*_*s*_ represents the total number of strong players, of which 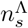play the focal strategy Λ. The probability mass function of the binomial distribution *B*(*k, n, p*) is taken as

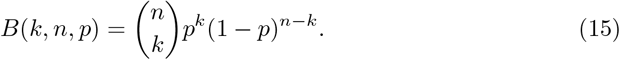

Similarly, counting for all possible combinations of the number of leaders 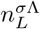with different strength levels *σ* and strategies Λ, the payoff turns out as

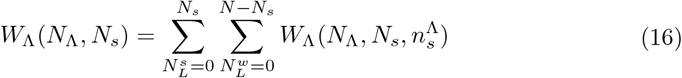

where 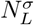represents the number of leaders with strength level *σ* and 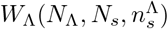 is defined as

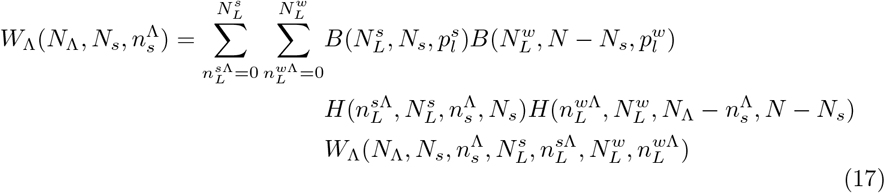

where 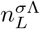represents the number of leaders with strength level *σ*, playing strategy Λ. The probability that an individual with strength level *σ* becomes a leader 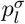 is calculated according to Eqs 3 or 7. The payoffs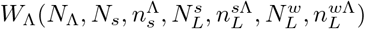are calculated following Eqs 10 and 11 taking into account 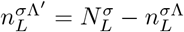

Once strategy payoffs are obtained, the transition probability between pairs of strategies is determined as fixation probabilities, i.e., the probability that a single mutant with a strategy *j* invades a population formed by *Z* − 1 individuals who follow a strategy *i* [48, 53, *54]:*

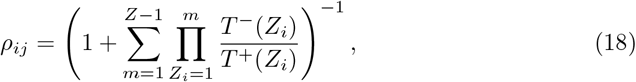

where *T* ^+^(*Z*_*i*_) is the probability that a resident individual imitates a mutant and *T*^*−*^(*Z*_*i*_) is the probability that a mutant imitates a resident one. Adopting a Fermi probability function [49], these imitation probabilities are given by

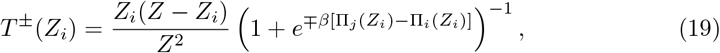

where *β* stands for the intensity of selection.

This approach allows us to use a reduced Markov chain to analyze the system [52, 53] and to compute the invasion diagram among all pairs of strategies, their stationary distributions, and an average level of cooperation across the different parameters of the model. Specifically, the non-diagonal elements of the transition matrix of the Markov chain are 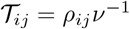, and the diagonal is *T*_*ii*_ = 1 *−* Σ _j_*T*_*ij*_, where *v* is the total number of strategies. The normalized eigenvector associated with the first eigenvalue of that matrix provides the stationary distribution *D*_*i*_, which represents the relative time the population spends adopting the strategy *i*. The transition probability 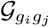 between two groups of strategies *g*_*i*_ and *g*_*j*_ is computed as:

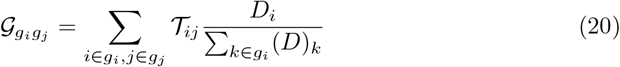

On the other hand, the average level of cooperation of each scenario is computed as

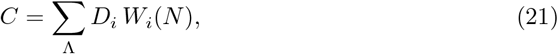

where the average payoff of individuals following strategy *i* in a homomorphic population is equivalent to the payoff *W*_*i*_(*N* ) they obtain playing within a group formed by them.

### Conditions for the emergence of cooperation under extreme heterogeneity

In order to assess the conditions for leadership to promote cooperation, as a first approach, we examine whether a cooperator mutant can invade a resident population of defectors and vice versa, in the regime when Δ_*l*_ and Δ_*f*_ are high. In this situation, strong individuals always become leaders and weak individuals always follow a strong individual (unless there is none in the group), i.e., 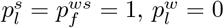, and 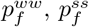 and 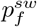 do not play any role. We also assume that the population is large enough for residents to obtain their average payoff through interactions within groups formed only by other residents, whereas the mutant is always interacting in a group formed by N-1 residents. In order to facilitate the analytical development and final expression, we consider *ϵ* = 0 in this very first approach. The difference between the average payoffs of the mutant and residents Π^*mut*^ − Π^*res*^ is positive if a mutant can invade the population and negative otherwise.

We first analyze this dynamics for **AllC** and **AllD** strategies. The average payoff of **AllD** and **AllC** resident individuals are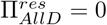and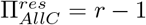 respectively. In order to deduce the payoff of a mutant individuals, we need first to establish the probabilities that cooperators and defectors cooperate or defect in a group formed by N-1 residents and a mutant. Applying Eq 1, it will be convenient to write the mutant payoffs as

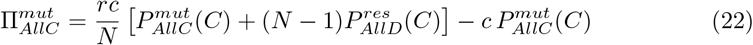

and

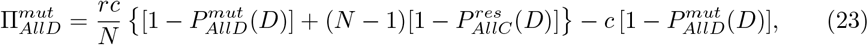

where 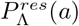and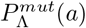 stand for the probabilities that an individual following strategy Λplays *a* =*{ C, D}* when resident or mutant, respectively. In this kind of groups, a mutant **AllC** player cooperates if it becomes a leader or if there are no leaders in the group; a **AllD** resident player, on the contrary, only cooperates when weak and follows a mutant strong **AllC** individual (among all the strong individuals in the group). The same reasoning can be applied for mutant **AllD** and resident **AllC** individuals but for the probability of defection. Thus, we obtain

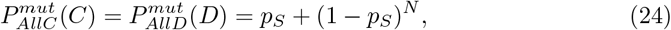

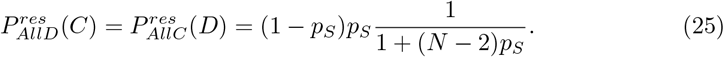

Therefore, the difference in payoffs between mutants and residents becomes

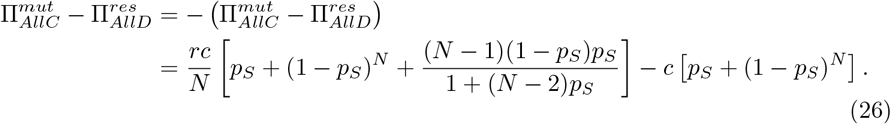

This relationship reveals that when **AllC** can invade **AllD, AllD** cannot invade **AllC**, and vice versa. It can also be easily seen that when there are no strong individuals (*p*_*S*_ = 0) or all individuals are strong (*p*_*S*_ = 1), i.e., when leadership is not in play, **AllC** dominates **AllD** if *r≥N*, which is the general condition for cooperation to prevail in the absence of any other mechanism as leadership. To analytically determine the probability of being strong (or the fraction of strong individuals in the population) for which **AllC** dominates over **AllD**, we need to simplify Eq 26 assuming that (1 *− p*_*S*_)^*N*^ *≈* 0. Since this approximation holds unless *p*_*S*_ is very small, it is possible to derive an upper bound for *p*_*S*_, but not a lower bound, which would be close to zero:

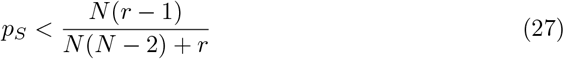

This analytical expression, even an approximate one, will be useful for interpreting further numerical results, since it relates the fraction of strong individuals, the natural/economic conditions, and the interacting group size. At first sight, it reveals that the higher *r* and the smaller the groups, the better for cooperation, which fully dominates only if *r ≥ N* .

Under the regime of high Δ_*l*_ and Δ_*f*_, 2-bit strategies **WDSC** and **NDLC** behave similarly to **AllC**, and **WCSD** and **NCLD** similarly to **AllD**, since the only differences lie in the first bit, i.e., how they act when they do not follow any leader and are weak or non-leading (which are equivalent in this regime). This occurs only with a probability (1*−p*_*S*_)^*N*^, which was neglected to deduce Eq 9. On the other hand, 4-bit strategies can be reduced to 2-bit ones since weak leaders and strong non-leaders are not possible under this regime. Therefore, there exist two main groups of strategies: **AllC**, [∗1], and [∗∗∗1] that promote cooperation when Eq 9 is satisfied and **AllD**, [∗0], and [∗∗∗0] that dominate otherwise. Following the same process described above, it can be easily deduced that **WDSC** and **NDLC** prevail over **AllC**, and **AllD** prevail over **WCSD** and **NCLD**, if (*N*−*r*)(1−*p*_*S*_)^*N*^ *>* 0, which is always satisfied provided that *r < N* . As a consequence, within strategy groups that promote cooperation or defection, individuals who defect when they are weak and/or not leading dominate those who cooperate in those situations.

When leadership is completely detached from strength (Δ = 0), it is possible to deduce analytically the conditions for cooperation to overcome complete defectors. In this situation, the probability of becoming a leader and the probability of an individual repeating the action of her leader are 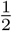 (given *f* = 0), i.e.,

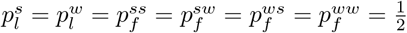 The probability of cooperating when playing **AllC, WDSC**, or **NDLC** strategies in a resident population formed by **AllD** individuals is, respectively:

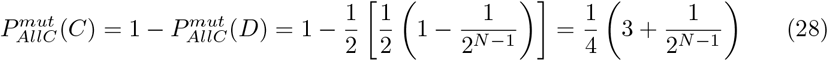

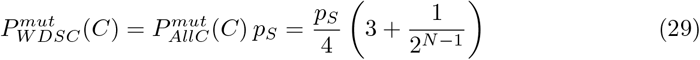

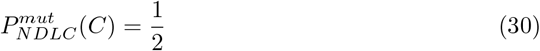

Assuming a large enough population and 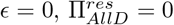and using Eq 1, we find that **AllC** and **WDSC** invade **AllD** if

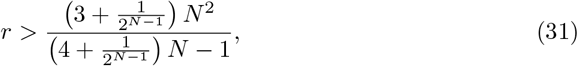

and **NDLC** does so if

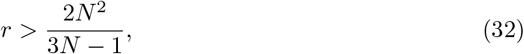

which for *N* = 9 correspond to *r >* 6.9 and *r >* 6.2, respectively. Therefore, when ecological conditions are favorable, cooperation is guaranteed if leaders and the decision to follow them are taken randomly.

## Supporting information

**S1 Cooperation levels in larger groups**. Cooperation levels as a function of *p*_*S*_ and Δ across the different strategy scenarios with *N* = 18 and *N* = 36 for the single-leader and multi-leader scenarios. The multiplication factor is set to *r* = 6*N/*9 to maintain consistency with the standard model for *N* = 9.

**S2 Cooperation levels for fixed** Δ_*f*_ **and varying** Δ_*l*_. Cooperation levels as a function of *p*_*S*_ and Δ_*l*_, with Δ_*f*_ = 1, for three representative values of the reward factor *r* across the different strategy scenarios.

**S3 Cooperation levels for fixed** Δ_*l*_ **and varying** Δ_*f*_ . Cooperation levels as a function of *p*_*S*_ and Δ_*f*_, with Δ_*l*_ = 1, for three representative values of the reward factor *r* across the different strategy scenarios.

**S4 Cooperation levels in the single leader scenarios**. Cooperation levels as a function of *p*_*S*_ and Δ for three representative values of the reward factor *r* across the different strategy scenarios.

**S5 Cooperation levels for increased error rate**. Cooperation levels as a function of *p*_*S*_ and Δ for *r* = 6, *ϵ* = 0.3 across the different strategy scenarios.

**S6 Stationary distribution for the S+L scenario**. Stationary distributions of all strategies in the S+L scenario and *r* = 6 as a function of *p*_*S*_ and Δ

**S7 Stationary distributions for the S and L scenarios**. Stationary distributions of all strategies in the S and L scenarios and *r* = 6 as a function of *p*_*S*_ and Δ

**S8 Invasion graphs of the main strategies in the S scenario**. Invasion graphs obtained for the S strategy scenario for representative values of *r, p*_*S*_, and Δ. All other parameter values and visualization criteria are the same as those used in Fig. 4.

**S9 Invasion graphs of the main strategies in the S+L scenario**. Invasion graphs obtained for the S+L strategy scenario for representative values of *r, p*_*S*_, and Δ. All other parameter values and visualization criteria are the same as those used in Fig. 4.

**S10 Invasion graphs of the main strategies in the L scenario**. Invasion graphs obtained for the L strategy scenario for representative values of *r, p*_*S*_, and Δ. All other parameter values and visualization criteria are the same as those used in Fig. 4.

**S11 Stationary distributions in the single-leader scenarios**. Stationary distributions of the dominant strategies for three representative values of the reward factor *r* across the parameter space defined by *p*_*S*_ and Δ for each strategy scenario.

